# Rational *in silico* discovery and serological validation of *Trypanosoma cruzi*-specific B-cell epitopes for high-precision Chagas disease diagnosis

**DOI:** 10.64898/2026.03.09.710567

**Authors:** Mayron Antonio Candia-Puma, Luis Daniel Goyzueta-Mamani, Haruna Luz Barazorda-Ccahuana, Raquel S.B. Câmara, Isabela A.G. Pereira, Ana L. Silva, Maíza M. Rodrigues, Bárbara P.N. Assis, Ana T. Chaves, Laís A.V.A. Corrêa, Manoel O. da Costa Rocha, Denise U. Gonçalves, Ana Alice Maia Gonçalves, Airam Barbosa de Moura, Alexsandro Sobreira Galdino, Ricardo Andrez Machado-de-Avila, Rodolfo Cordeiro Giunchetti, Eduardo Antonio Ferraz Coelho, Miguel Angel Chávez-Fumagalli

## Abstract

Chagas disease is caused by the parasite *Trypanosoma cruzi* and remains a neglected tropical disease presenting a substantial global health burden. Crude antigen-based assays have historically been limited in specificity; however, even contemporary recombinant-antigen tests may exhibit residual cross-reactivity, depending on antigen composition and geographic context. To overcome this limitation, this study developed a novel diagnostic strategy that integrates computational and experimental approaches to identify specific linear B-cell epitopes within the *T. cruzi* proteome. The strategy was developed to exclude sequences homologous to *H. sapiens* and *Leishmania* spp. proteins, thereby minimizing potential cross-reactivity. Using a consensus approach across five prediction algorithms, B-cell epitopes were identified and subsequently clustered to reveal conserved, immunoreactive consensus sequences. The peptide sequences were characterized for optimal physicochemical properties and subsequently modeled to interact with a human antibody using protein-peptide docking and molecular dynamics simulations to assess complex stability. The most promising candidates were chemically synthesized and validated using ELISA against a cohort comprising Chagas disease patients (chronic indeterminate and cardiac forms), healthy donors, and a cross-reactive control group (visceral and tegumentary leishmaniasis and leprosy). From the initial set of 19,245 proteins, the multi-tiered bioinformatic analysis identified 4,431 unique, non-homologous sequences. Consensus prediction yielded 401 high-confidence epitopes, which were refined to 179 structurally stable candidates. Computational analyses identified five top-ranking epitopes capable of forming high-affinity, stable complexes with a human antibody. Experimental validation confirmed the high diagnostic accuracy of two epitopes, which demonstrated exceptional diagnostic performance: Epitope 4 and Epitope 5 achieved 100% sensitivity. Notably, Epitope 5 exhibited superior specificity, reaching 96.67% against healthy controls and 90.91% against the cross-reactive group. This study establishes a basis for the development of an improved immunoassay for Chagas disease and provides a reproducible framework for targeted epitope discovery. Consequently, this study validates a high-precision computational pipeline capable of discovering *T. cruzi*-specific antigens that effectively circumvent cross-reactivity with *Leishmania* spp., proposing Epitope 5 as a qualified candidate for reliable serological diagnosis in co-endemic regions.

## Introduction

Chagas disease (CD), caused by the hemoflagellate parasite *Trypanosoma cruzi*, constitutes a neglected tropical disease of global impact with devastating consequences (1). Classified by the WHO as one of the deadliest parasitosis in the Americas, it affects an estimated 6–7 million people in endemic regions of Latin America, with 12,000 deaths each year—mainly from Chagasic cardiomyopathy, megacolon, and sudden death (2). Globalization has extended its presence to more than 30 non-endemic countries through migration and blood transfusion, transforming it into a transcontinental challenge (3). This epidemiological burden underscores the urgency of precise diagnostics, particularly during indeterminate chronic phases and in blood banks, where serological testing is indispensable.

In the complex landscape of CD, where the often-silent progression of pathology and low parasitemia render direct parasitological methods insufficient during the chronic stage, serology stands as the definitive bridge to timely patient management and epidemiological control (4). Consequently, the detection of specific antibodies is the cornerstone of CD diagnosis. The distinct roles of immunoglobulin isotypes (IgG, IgM, and IgA) provide critical insights into the stage and dynamics of the infection. IgM antibodies are the first responders, appearing during the acute phase, but their transient nature and lower titers make them less reliable for routine screening (5,6). In contrast, IgG antibodies emerge later and persist indefinitely throughout the chronic phase, making them the primary target for confirmatory serological testing in individuals with suspected chronic infection and for blood bank screening (7). Notably, the detection of IgA, though less routinely employed, has shown promise as a potential marker of active infection and therapeutic response, as its levels may decline more rapidly after successful treatment compared to total IgG (8,9). Therefore, a comprehensive diagnostic tool should ideally account for these isotypes to maximize clinical utility across different phases of the disease.

Nevertheless, the current diagnostic standard—two serological assays (ELISA, HAI, IIF) employing crude (10) or semi-purified (11) antigens—suffers critical limitations that compromise reliability. Most notably, low specificity arises from cross-reactivity with *Leishmania* spp. (especially *L. braziliensis* and *L. infantum*), syphilis and autoimmune diseases, yielding up to 15 % false-positive results in co-endemic areas (4,5). Additional issues include reduced sensitivity in immunocompromised individuals, lack of standardization among commercial kits, and failure to discriminate active from resolved infection (5,12,13). These shortcomings carry serious clinical (erroneous treatment, patient anxiety) and operational (discarded blood units, elevated costs) consequences, highlighting the pressing need for high-precision diagnostic alternatives.

In this context, chemically synthesized soluble linear B-cell epitopes emerge as a promising solution. These short peptides (< 20 amino acids), corresponding to specific antigenic regions recognized by antibodies, enable the exclusion of non-specific immunogenic components present in crude extracts, the principal source of cross-reactivity (14). Their chemical synthesis facilitates the development of highly standardized, reproducible, and scalable assays, with the potential to rationally optimize both sensitivity and specificity (15,16). Crucially, synthetic peptides are inherently stable and do not require a cold chain for storage and transport (17,18). This attribute represents a significant logistical advantage for the deployment of diagnostic tests in remote, rural areas (where Chagas disease is predominantly endemic) and where infrastructure, including reliable electricity for refrigeration, is often limited or unavailable (19).

Traditional epitope identification via immunological screening, however, is inherently slow, costly, and low-throughput (20). To circumvent these limitations, *in silico* strategies offer a robust alternative. Bioinformatic tools (such as BepiPred, ABCPred, and APP) facilitate the rapid and cost-effective prediction of candidate epitopes directly from protein sequences (21,22). Yet, while previous studies have applied these computational approaches to Chagas disease (23,24), a critical gap remains: the identification of epitopes with exclusive specificity for *T. cruzi*. Prior efforts have struggled to effectively filter out conserved antigenic regions shared with *Leishmania* spp. and other phylogenetically related pathogens, leading to persistent cross-reactivity (4). This limitation is particularly detrimental in co-endemic regions of South America, where the high degree of sequence homology between these parasites makes serological discrimination between Chagas disease and leishmaniasis a formidable challenge.

This study proposes an integrated *in silico*–experimental approach to identify and validate a *T. cruzi*-specific linear B-cell epitope. The central hypothesis suggests that rigorous bioinformatic design, focused on excluding homology with the proteomes of *Leishmania* spp. and *Homo sapiens*, can yield peptides that, when implemented in ELISA, achieve high sensitivity and specificity, thereby eliminating false positive results. By closing this critical gap, the study aims to provide a transformative diagnostic tool for epidemiological control. Therefore, this study aimed to identify and experimentally validate species-specific linear B-cell epitopes from *T. cruzi* for diagnostic application.

## Material and Methods

### *In silico* prediction of *T. cruzi*-specific B-cell epitopes

#### Obtaining and initial processing of the reference proteome

The reference proteome of *T. cruzi* was obtained from the UniProt database (http://www.uniprot.org, accessed on 15 March 2023) (25), prioritizing a high-coverage, non-redundant protein set. Initial physicochemical characterization—including molecular weight (kDa) and isoelectric point (pI)—was performed for all proteins using the Compute pI/Mw tool on the ExPASy server (https://web.expasy.org/compute_pi/, accessed on 20 March 2023) (26).

#### Filtering by homology with host and related parasites

To diminish potential cross-reactivity in diagnostic applications, sequences homologous to *H. sapiens* (taxid:9606) and *Leishmania* (taxid:5658) proteins were excluded. Each protein in the reference proteome underwent pairwise comparison against both host and parasite databases using BLASTp (https://blast.ncbi.nlm.nih.gov/Blast.cgi, accessed on 14 April 2023) (27). Homology was defined by stringent thresholds: an e-value ≤ 0.005 and/or a bit score ≥ 100.0. Consequently, sequences showing homology were excluded, while those lacking significant similarity were retained for downstream analyses to ensure candidate epitopes are uniquely recognizable as *T. cruzi*-specific antigens.

#### Prediction of linear B cell epitopes

Linear B-cell epitope prediction was performed using five algorithms: AAP (http://ailab-projects2.ist.psu.edu/bcpred/predict.html, last accessed on 24 February 2024) (28,29), ABCpred (https://webs.iiitd.edu.in/raghava/abcpred/ABC_submission.html, last accessed on 24 February 2024) (30–32), BCpred (http://ailab-projects2.ist.psu.edu/bcpred/predict.html, last accessed on 24 February 2024) (30,32,33), BepiPred-2.0 (https://services.healthtech.dtu.dk/services/BepiPred-2.0/, last accessed on 24 February 2024) (34–36), and FBCPred (http://ailab-projects2.ist.psu.edu/bcpred/predict.html, last accessed on 24 February 2024) (30,32,37). These tools incorporate distinct computational approaches—support vector machines, recurrent neural networks, string kernel methods, random forests, and subsequence kernel methods, respectively—to identify epitopes with a high likelihood of efficient processing by B cells. A specific prediction score threshold was applied for each algorithm, selected to optimize specificity and ensure high-confidence selections. The exact threshold values used are detailed in Table 1.

**Table 1.** Linear predictors of B-cell epitopes.

| Tool | Method | Prediction threshold | Specificity | Epitope size | Web Link | References |
| --- | --- | --- | --- | --- | --- | --- |
| AAP | Support vector machine + amino acid pair antigenicity scale | 0.70 | 75% | 12 | http://ailab-projects2.ist.psu.edu/bcpr ed/predict.html | (28,29) |
| BCpred | Support vector machine + k-mer subsequence kernel | 0.80 | 75% | 12 | http://ailab-projects2.ist.psu.edu/bcpr ed/predict.html | (30,32,33) |
| FBCPred | Support vector machine + flexible subsequence kernel | 0.85 | 75% | 12 | http://ailab-projects2.ist.psu.edu/bcpr ed/predict.html | (30,32,37) |
| ABCpred | Recurrent neural network (Jordan network) | 0.80 | 95.50% | 12 | https://webs.iitd.edu.in/raghava/abcpr ed/ABC_submission.html | (30–32) |
| BepiPred-2.0 | Random forest + physicochemical features | 0.55 | 81.66% | ≥ 12 | https://services.healthtech.dtu.dk/services/BepiPred-2.0/ | (34–36) |

#### Identification of consensus epitopes

To identify candidate epitopes with a higher likelihood of being immunoreactive and thus prioritize them for the development of a diagnostic test, the resulting epitopes were compared, and consensus epitopes were determined using the clustering tool for similar sequences from the Immune Epitope Database (IEDB) (http://tools.iedb.org/cluster/, last accessed on 24 August 2024) (38). This consensus approach is based on its ability to rank candidates according to their conservation and likelihood of immune response. The parameters used included a minimum sequence identity threshold of 70% and a peptide length ranging from 12 to 16 amino acids (39,40).

#### Analysis of physicochemical properties

The physicochemical properties of the selected sequences were analyzed using the ProtParam tool on the ExPASy server (https://web.expasy.org/protparam/, last accessed on 27 September 2024) (26) to determine the following parameters: (1) amino acid composition; (2) theoretical pI calculated using the Bjellqvist method; (3) stability predictors, including the instability index (values >40 indicating instability) and aliphatic index (thermostability proxy); and (4) hydrophobicity via the grand average of hydropathicity (GRAVY; positive values denote hydrophobic character).

#### Retrieval of structural, sequential data and delineation of the paratope

A search was conducted in the Protein Data Bank (PDB) (https://www.rcsb.org/) to identify three-dimensional entries of human immunoglobulins from the IgA (3M8O) (41), IgG (1MIM) (42), and IgM (1DN0) (43) classes. For each structure, the .pdb files and FASTA sequences of the heavy and light chains were downloaded simultaneously, ensuring that primary and structural data were linked from the outset of the analysis. If an entry contained missing regions or incomplete domains, these portions were reconstructed through homology modeling using the SWISS-MODEL server (https://swissmodel.expasy.org/, last accessed on 01 October 2024) (44), generating complete models with appropriate confidence levels. The stereochemical quality and backbone conformation integrity of all initial templates and the subsequently generated homology models were rigorously validated. This was achieved by analyzing the phi (φ) and psi (ψ) dihedral angle distributions via Ramachandran plots computed using the PROCHECK software suite (45). Given the presence of flexible linker regions and loops characteristic of immunoglobulin domains, a model was considered acceptable for subsequent analysis if a minimum of 90% of all non-glycine, non-proline residues were located within the most favored and additionally allowed regions of the conformational space (46). All refined models met or exceeded this threshold, ensuring the structural plausibility of the rebuilt domains for our comparative analysis. The heavy and light chains were analyzed in parallel using IMGT/DomainGapAlign (https://www.imgt.org/3Dstructure-DB/cgi/DomainGapAlign.cgi, accessed on 04 October 2024), Parapred (https://www-cohsoftware.ch.cam.ac.uk/index.php/Parapred, accessed on 04 October 2024), and LYRA (http://tools.iedb.org/lyra/, accessed on 04 October 2024), three complementary algorithms for predicting antibody-antigen contact regions (47–49). Overlapping the results enabled the construction of a consensus paratope sequence, minimizing tool-specific biases and enhancing the reliability of identifying critical residues for antigen recognition.

#### Epitope modeling

Epitopes deemed stable in the previous stage underwent de novo three-dimensional prediction using the PEP-FOLD 3.5 server (https://mobyle.rpbs.univ-paris-diderot.fr/cgi-bin/portal.py#forms::PEP-FOLD3, accessed on 09 October 2024) (50). For each linear sequence, conformational sampling was performed utilizing the coarse-grained sOPEP (single-point optimized potential for efficient peptide structure prediction) force field, with the default parameters recommended for peptides spanning 5 to 50 residues. The primary quantitative metric for evaluating the energetic quality and conformational stability of each generated model was the sOPEP Energy. This score serves as the key thermodynamic indicator of modeling efficiency within the PEP-FOLD framework, where a lower, more negative value correlates with a more stable, energetically favorable conformation, reflecting a higher likelihood of representing a native-like fold in solution (51).

#### Protein-peptide molecular docking and molecular dynamics

Selected Fab fragments and epitopes were docked with HPEPDOCK 2.0 (http://huanglab.phys.hust.edu.cn/hpepdock, last accessed on 16 October 2024) and HawkDock (http://cadd.zju.edu.cn/hawkdock/, last accessed on 16 October 2024), engines integrating peptide flexibility and free energy estimation (52,53). Top-scoring complexes were visually inspected in UCSF ChimeraX to validate binding geometries consistent with the paratope and rule out computational artifacts. To prioritize computational resources, only complexes with a docking score exceeding the mean plus one standard deviation in both programs were retained (a statistical outlier threshold that, under a normal distribution, targets the upper ∼16% of results from each independent screen) (54). This threshold balanced sensitivity and specificity, focusing simulations on interactions with a higher likelihood of biological relevance.

Molecular dynamics simulations of the protein-peptide complexes identified through docking were performed using Schrödinger Suite 2025-2, specifically the DESMOND module (55). Protein systems were prepared and solvated in a TIP3P water model within a cubic periodic boundary box, neutralized with 0.15 M NaCl under the OPLS5 force field. Energy minimization (100 ps) preceded production MD simulations of 500 ns under NPT ensemble conditions at 300 K and 1.01325 bar, with trajectory frames recorded every 500 ps (yielding 1000 frames). Post-simulation analysis utilized Schrödinger’s Simulation Interaction Diagram tool to quantify root mean square deviation (RMSD), percentage of structural stability energy (%SSE), and residue index.

#### Serological evaluation of candidate epitopes against sera from patients Chemical synthesis of peptides

Candidate epitopes, selected based on in silico screening for immunogenicity and specificity, were synthesized using standard Fmoc (9-fluorenylmethoxycarbonyl) solid-phase peptide synthesis (SPPS) on a polystyrene resin, following previously described protocols (56–59). The process was meticulously executed through iterative cycles of N-terminal Fmoc deprotection with 25% (v/v) 4-methyl-piperidine in dimethylformamide (DMF), followed by extensive washing with DMF and dichloromethane to remove reaction byproducts. Subsequent coupling of each Fmoc-protected amino acid was qualitatively verified after each cycle using the bromophenol blue test to ensure reaction completion. Following the assembly of the full sequence, the nascent peptide was simultaneously cleaved from the resin and globally deprotected via treatment with a chilled cleavage cocktail—a precise mixture of trifluoroacetic acid (TFA; 95% v/v), triethylsilane (TES; 2.5% v/v) as a carbocation scavenger, and ultrapure water (2.5% v/v)—for 3 hours under gentle agitation. The crude peptide was then precipitated in a copious volume of cold diethyl ether and incubated at -20°C for 18 hours to maximize yield. The resulting pellet was isolated by centrifugation at 4500×g for 10 minutes at 4°C, subjected to repeated ether washes to purge organic impurities, and finally lyophilized to obtain the pure product as a stable powder, which was stored at -20°C and reconstituted in Milli-Q water at 1.0 mg/mL immediately before use. Five peptides were synthesized and named epitope 1 to 5.

#### Ethical statement and Study population

The study was approved by the Ethics Committee of the Federal University of Minas Gerais (UFMG, Belo Horizonte, Minas Gerais, Brazil), logged under protocol number CAAE-32343114.9.0000.5149. All participants were adults and signed the informed consent form at the moment of inclusion.

A total of 178 subjects were enrolled in the study. Blood was obtained through venipuncture using a sterile 20 mL vacuum collection tube with clot activator and separator gel. Subsequently, tubes were centrifuged at 3,500 x *g* for 15 min at 4°C, and the serum was separated and stored at -70°C until use. A total of 88 serum samples were obtained from Chagas disease patients from the Center for Training and Reference in Infectious and Parasitic Diseases (CTR DIP-Orestes Diniz) in Belo Horizonte, Minas Gerais, Brazil. Patients are presented with an indeterminate form (n = 43) or Chagas cardiomyopathy (n = 45) clinical disease. Parasitological confirmation of *T. cruzi* infection was performed by hemoculture in combination with specific ELISA or indirect hemagglutination assay (IHA). Leishmaniasis was diagnosed by clinical evaluation and parasitological test for tegumentary leishmaniasis (TL) (Giemsa-stained smears from lesion fragments and PCR to detect *L. braziliensis* kDNA) and visceral leishmaniasis (VL) (aspirates from spleen and/or bone marrow and PCR to detect *L. infantum* kDNA) (n=20). TL was clinically stratified in cutaneous (n=10) and mucosal (n=10) leishmaniasis. Leprosy was diagnosed by clinical evaluation, slit-skin smears, and histopathological examination of the lesion biopsy. Patients were classified according to WHO criteria in presenting either multibacillary (MB, n=10) or paucibacillary (PB, n=10) form. Sera samples from healthy subjects living in the endemic region of disease, Poços de Caldas, in the state of Minas Gerais (n=30), who did not present clinical signs or symptoms of any infectious disease at the moment of collection, were also used.

#### ELISA

Previous titration curves were performed to determine the most appropriate concentration of antigens and antibody/sample dilutions to be used in the ELISA experiments. Briefly, 96-well high-binding plates (Thermo Fisher, USA) were coated with each synthetic peptide (4 µg, 6 µg, 4 µg, and 8 µg per well, respectively), which were diluted in carbonate buffer pH 9.6 and incubated for 1 h at 37°C. Wells were blocked with 200 µL of Phosphate Buffered Saline (PBS 1x pH 7.4) plus Tween 20 (PBS-T) containing 2% (w/s) casein for 1 h at 37°C. Plates were washed three times with PBS-T and incubated with 50 µL/well of serum samples, which were diluted 1:100 in PBS-T containing 2% casein, for 1 h at 37°C. After washing plates three times with PBS-T, 100 µL/well of peroxidase-conjugated anti-human IgG antibody (A1811, Invitrogen, USA), which was diluted 1:5,000 in PBB-T plus 2% casein, was added to the wells, and plates were incubated for 1 h at 37°C. Next, wells were washed three times with PBS-T, and reactions were developed by adding 3,3‘,5,5’; tetramethylbenzidine (TMB, Scienco, Brazil) at room temperature for 20 min and in the dark. Reactions were stopped by adding 50 µL of 2N H_2_SO_4_, and the optical density (OD) values were read in a microplate spectrophotometer (Molecular Devices, Spectra Max Plus, Canada), at 450 nm (60).

#### Statistical analysis

Results were entered into Microsoft Excel (version 10.0) spreadsheets and analyzed with GraphPad Prism version 10.0.2 for Windows (GraphPad Software, USA, www.graphpad.com). The cut-off values were calculated through Receiver Operating Characteristic (ROC) curves. The ROC curves were plotted with the values from samples from *T. cruzi*-infected patients versus those from control groups. For cutoff selection, we use the highest positive likelihood ratio (LR+), defined as the ratio of the probability of a positive test in patients to that in controls. Tables of contingency and Fisher’s exact test (P < 0.05) were used to calculate: Sensitivity (Se), Specificity (Sp), and the 95 % Confidence Interval (95 % CI). An ordinary one-way analysis of variance (ANOVA) followed by Tukey’s multiple comparison test, which assumes normal Gaussian distribution and a single pooled variance, was used for group comparisons. Differences were considered significant with *P* < 0.05.

## Results

### Comprehensive rational design pipeline for epitope discovery

To systematically identify and validate novel diagnostic markers for Chagas disease, we developed a sequential, ten-step computational and experimental framework, as illustrated in Figure 1. The pipeline initiates with Reference Proteome Retrieval from UniProt (Step 1), followed by rigorous Cross-Reactivity Filtering (Step 2) to eliminate homologous sequences. Candidate sequences are then subjected to Linear Epitope Prediction (Step 3) using multiple algorithms, culminating in a Consensus Identification (Step 4) to minimize false positives. The selected epitopes undergo Physicochemical Analysis (Step 5) to ensure structural stability. Promising candidates advance to De Novo Modeling (Step 6) and subsequent Molecular Docking and Dynamics (Step 7) to evaluate their theoretical binding affinity and stability with human immunoglobulins. Successfully prioritized targets are then produced via Chemical Synthesis (Step 8) and subjected to Serological Evaluation (Step 9) using patient cohorts. Finally, the diagnostic performance of the assays is rigorously evaluated through Statistical Analysis (Step 10).

**Figure 1.**
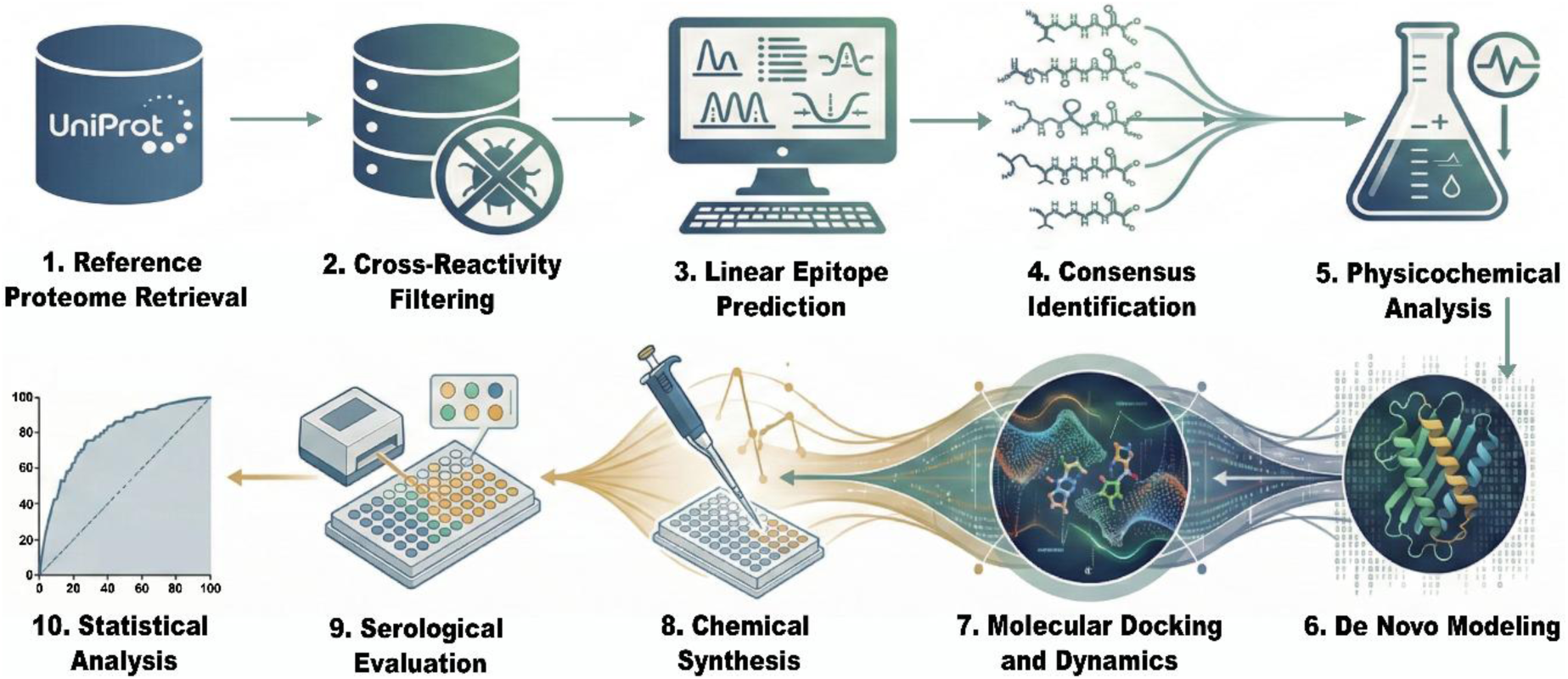
Integrative immunoinformatics and experimental pipeline for the discovery and validation of Trypanosoma cruzi B-cell epitopes.

### *In silico* identification of antigenic epitopes

#### Identification, screening, and physicochemical characterization of non-homologous B-cell epitopes

The *T. cruzi* reference proteome (strain CL Brener - UniProt ID: 353153) contains 19,245 entries. We applied a quality control filter excluding 3,069 (15.9%) truncated protein fragments, as they impede accurate physicochemical parameter calculation (Fig. 2A). To focus our analysis on the uncharacterized proteome, we further excluded the 49 (0.3%) manually reviewed in UniProt (Fig. 2B). The final dataset comprises 16,127 unreviewed, full-length proteins (Fig. 2C).

**Fig. 2.**
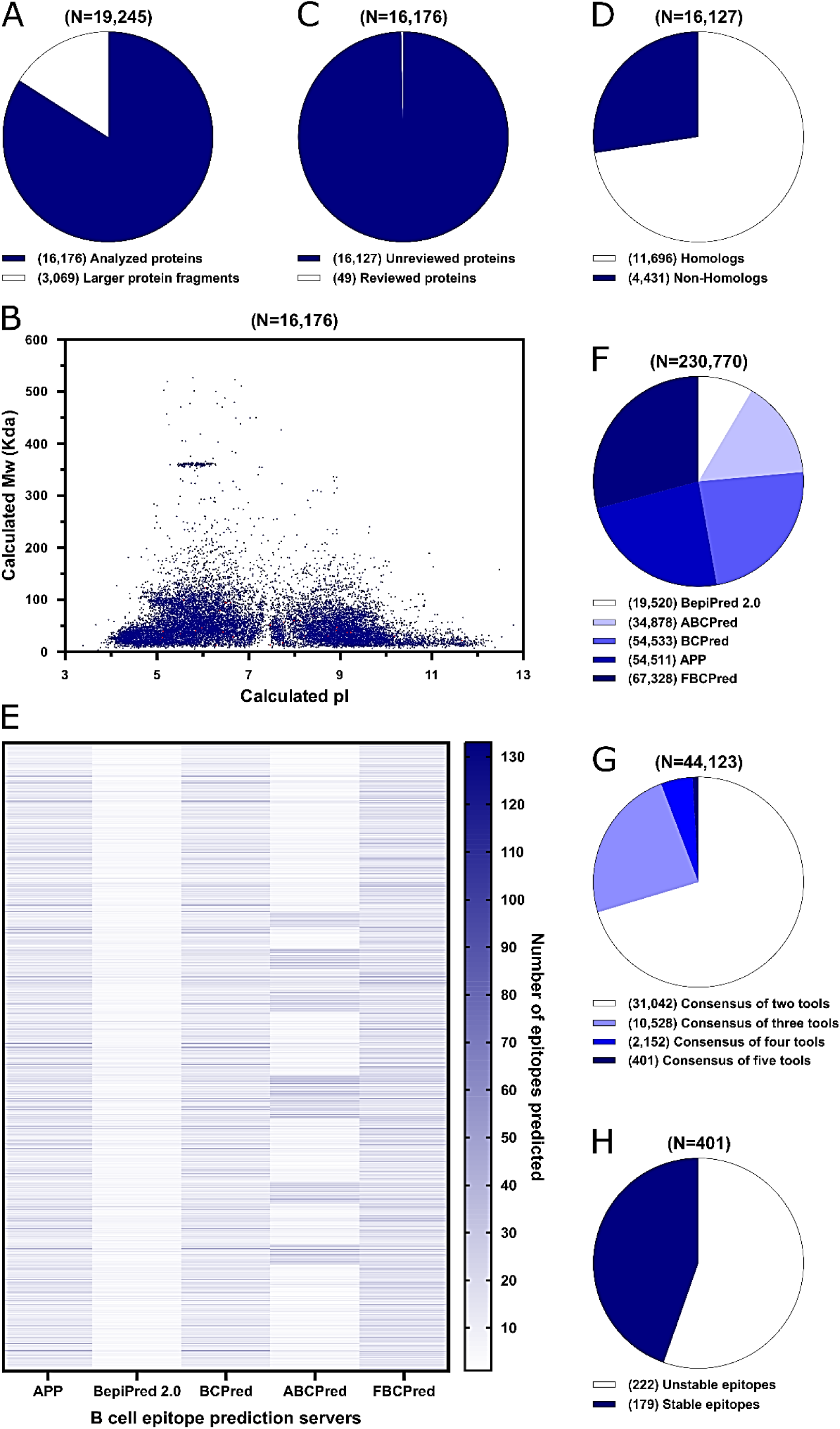
I*n silico* analysis of B-cell epitopes in the *Trypanosoma cruzi* proteome. (A) Total proteome (19,245 proteins) excluding truncated fragments (3,069). (B) The calculated molecular weight (Mw) and isoelectric point (pI) of 16,176 proteins are provided. The status of the proteins is as follows: unreviewed (blue dots - 16,127) and reviewed (red dots - 49). (C) Selection of unreviewed full-length proteins (16,127) after excluding reviewed ones (49). (D) Identification of 4,431 non-homologous proteins (H. sapiens/Leishmania). (E) Heatmap of predicted epitopes for non-homologous proteins using five B-cell epitope prediction servers (APP, BepiPred 2.0, BCPred, ABCPred, FBCPred). (F) Prediction of B-cell epitopes in non-homologous proteins using five servers (counts: APP=54,511, BepiPred=19,520, BCPred=54,533, ABCPred=34,878, FBCPred=67,328). (G) Consensus epitopes: 31,042 (2 tools), 10,528 (3 tools), 2,152 (4 tools), 401 (5 tools). (H) 179/401 total consensus epitopes showed structural stability (ProtParam).

Sequence analysis using BLASTp identified 113 sequences with an expected e-value greater than 0.005 and a bit score less than 100.0 against reference proteomes of *H. sapiens* and/or *Leishmania*. Additionally, 4,318 sequences showed no significant similarity. In total, 4,431 sequences (Fig. 2D) did not exhibit detectable homology with proteins from these organisms, suggesting the presence of unique or highly specific elements in *T. cruzi*.

To visualize potential epitopes on non-homologous proteins, a heatmap was generated using five specialized B-cell epitope prediction servers: APP, BepiPred 2.0, BCPred, ABCPred, and FBCPred. This diverse strategy, which employs multiple algorithms, enables a comprehensive assessment of regions likely to be recognized by the immune system, accounting for the structural and sequence variability of the target proteins. The heatmap provides an intuitive visual representation of the distribution and intensity of the predicted epitopes (Fig. 2E). The total number of epitopes predicted by each server was: APP identified 54,511, BepiPred 2.0 identified 19,520, BCPred identified 54,533, ABCPred identified 34,878, and FBCPred identified 67,328 (Fig. 2F). This substantial variability in counts reflects the inherent differences in the algorithms and methods employed by each tool, highlighting how the choice of method significantly influences the identified immunogenic landscape (61,62).

To increase confidence in the predictions, a consensus analysis was performed across the five servers. 31,042 epitopes were identified with agreement across at least two tools. The consensus across three tools yielded 10,528 epitopes, the consensus across four tools yielded 2,152 epitopes, and the highest consensus (five tools) identified 401 epitopes (Fig. 2G). This approach stratifies predictions based on the level of agreement, providing a confidence gradient where higher consensus reduces the likelihood of false positives.

Of the 401 epitopes predicted by consensus across the five tools, 179 exhibited stabilities according to the instability index calculated using the ProtParam tool on the ExPASy server (Fig. 2H). This predicted structural stability is a crucial attribute for the potential biological functionality of these epitopes, as it increases the likelihood that they will maintain a conformation recognizable by the immune system (63,64).

### Structural modeling and validation of epitope-paratope interfaces

Representative human Fabs of each isotype were selected, strictly complying with intact domains, absence of mutations, and human origin, and their linked .pdb files and FASTA sequences were downloaded. The IgA Fab (PDB 3M8O, 1.55 Å) stood out, with R_work 0.153, R_free 0.182, 0% Ramachandran outliers, clash score 9, and only 1.3% side-chain deviations; the IgG Fab (PDB 1MIM, 2.60 Å) showed moderate quality (R_work 0.196, clash score 10, 2.4% outliers, and 12.1% misplaced side chains); and the IgM Fab (PDB 1DN0, 2.28 Å) exhibited similar metrics (R_work 0.180, R_free 0.240, clash score 22, 1.2% outliers, and 12.7% misplaced side chains). Missing regions in any of these entries were completed through homology modeling in SWISS-MODEL, generating comprehensive models with suitable confidence levels before proceeding with further comparative analyses.

The combination of IMGT/DomainGapAlign, Parapred, and LYRA converged on a consensus paratope for each chain, revealing distinct CDR profiles across isotypes: IgA heavy chain (H-CDR1: ASGFTLSGSNV, H-CDR2: RIKRNAESDATA, H-CDR3: VIRGDVYNRQWG; light chain L-CDR1: SCRSSQSLLRRDGHNDLEWY, L-CDR2: IYLGSTRASGV, L-CDR3: YCMQNKQTPLTFG), IgG (H-CDR1: ASGYSFTRYWM, H-CDR2: AIYPGNSDTS, H-CDR3: SRDYGYYFDFWG; L-CDR1: TCSASSSRSYMQWY, L-CDR2: IYDTSKLASGV, L-CDR3: YCHQRSSYTFG), and IgM (H-CDR1: VYGGSFSDYYW, H-CDR2: EINHSGSTN, H-CDR3: ARPPHDTSGHYWNYWG; L-CDR1: SCGASQSVSSNYLAWY, L-CDR2: IYDASSRATGI, L-CDR3: YCQQYGSSPLTFG). Their distribution, visualized in pearl-necklace plots, showed contact regions as densely clustered beads along Fab sequences (Fig. 3). Consistently, the heavy chain paratopes spanned longer segments than those of the light chains, underscoring their dominant role in antigen recognition. This representation facilitated inter-isotype comparison and revealed conserved patterns that may underpin the broad functional versatility of immunoglobulins.

**Fig. 3.**
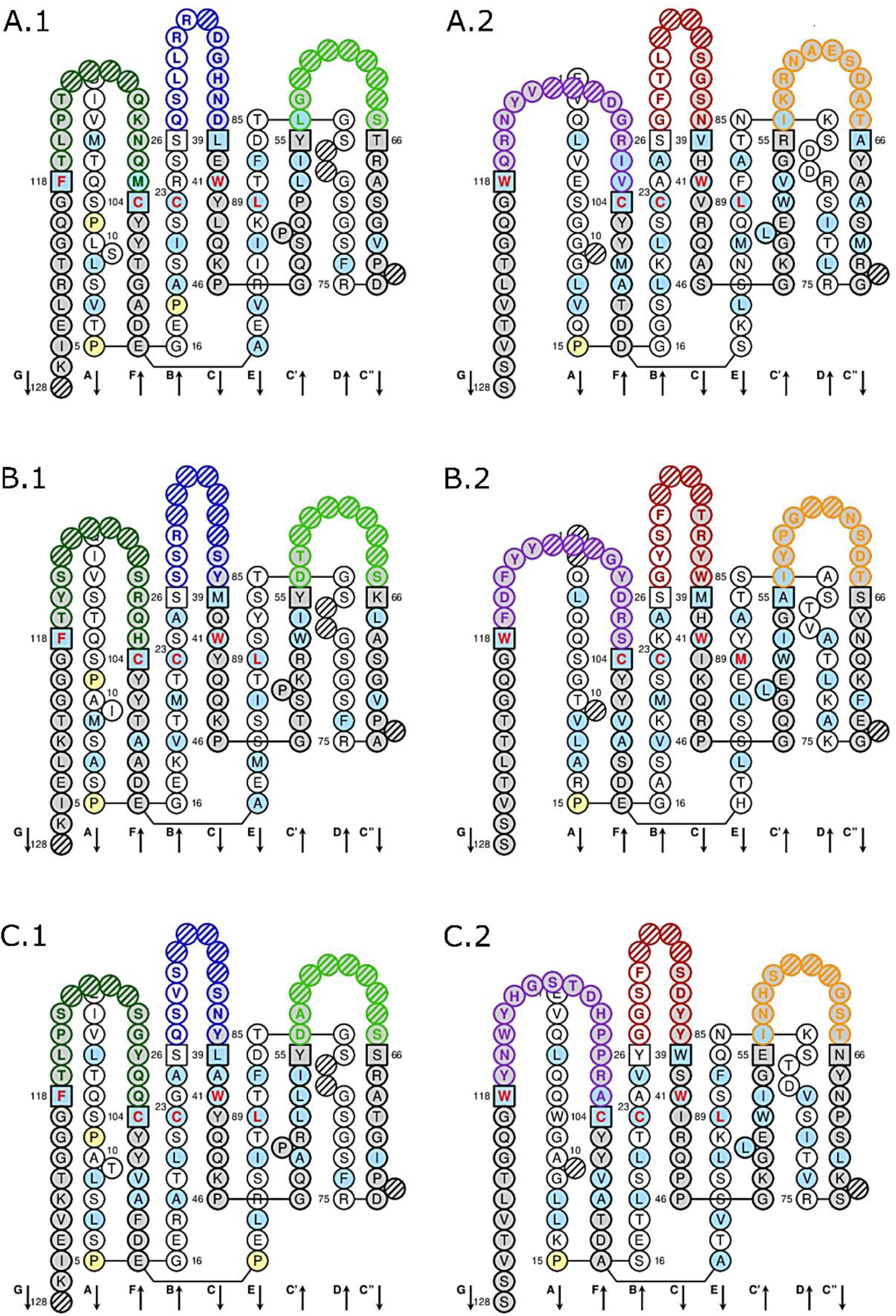
Pearl necklace plot representation of the paratopes in the Fab regions of human immunoglobulins across three isotypes (IgA, IgG, IgM), as determined by the IMGT/DomainGapAlign tool. a) IgA heavy chain, b) IgA light chain, c) IgG heavy chain, d) IgG light chain, e) IgM heavy chain, f) IgM light chain.

The 179 epitopes obtained from the *in silico* analysis of the *T. cruzi* proteome were modeled using PEP-FOLD 3.5, preserving structural motifs compatible with antibody binding. Their compactness and lack of disordered regions favored the subsequent docking phase. The three-dimensional prediction allowed preliminary inspection of key residue accessibility and the elimination of sterically unviable conformations, ensuring that only biologically relevant structures advanced to immunological recognition assessment via molecular docking.

### Molecular Docking and Dynamic Stability Analysis of the Immune Complex

Parallel docking with HPEPDOCK 2.0 and HawkDock revealed high-affinity interactions selective for each immunoglobulin class. In total, 1 epitope was identified for IgA (HPEPDOCK: -216.46; HawkDock: -3912.81), 1 for IgM (HPEPDOCK: -220.14; HawkDock: -3421.17), and 3 for IgG (Ep2 HPEPDOCK: - 240.53, HawkDock: -3907.48; Ep3 HPEPDOCK: -211.97, HawkDock: -3607.84; Ep4 HPEPDOCK: -227.15, HawkDock: -4260.45), all with scores above the mean + 1 standard deviation in both tools (Table 2). Visualizations in UCSF ChimeraX confirmed consistent binding geometries: the epitopes nestled into the cavities outlined by the paratopes, forming complementary hydrogen bond networks and hydrophobic contacts (Fig. 4).

**Fig. 4.**
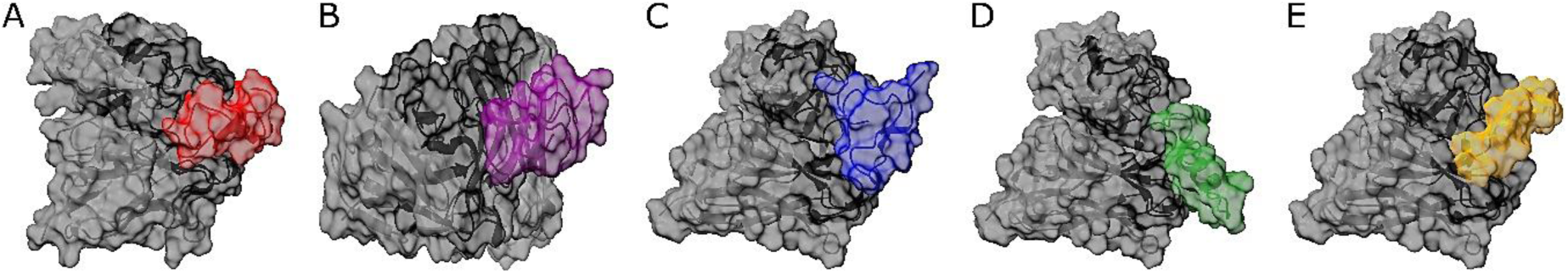
Molecular docking models depicting epitope binding to immunoglobulin FAB regions: (A) Epitope bound to IgA, (B) Epitope bound to IgM, and (C–E) Distinct epitopes bound to IgG. The FAB region is highlighted in gray, the paratope in black, and epitope colors (A–E) correspond to the sequences specified in Table 2.

**Table 2.**
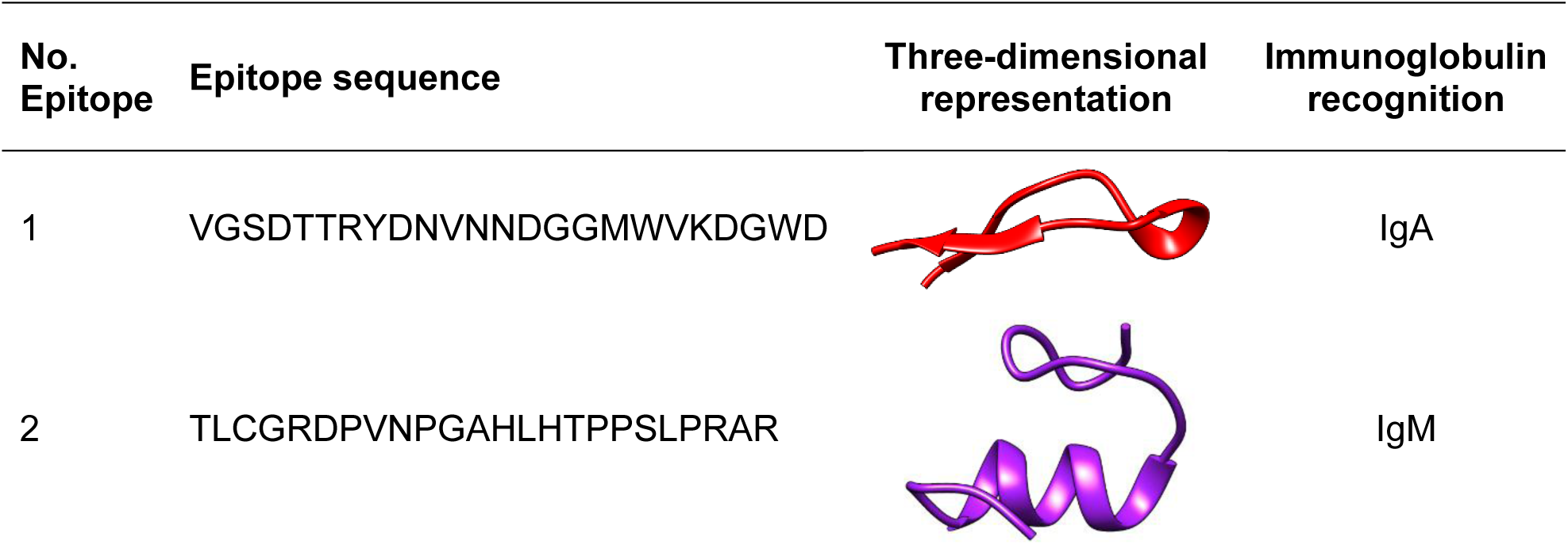
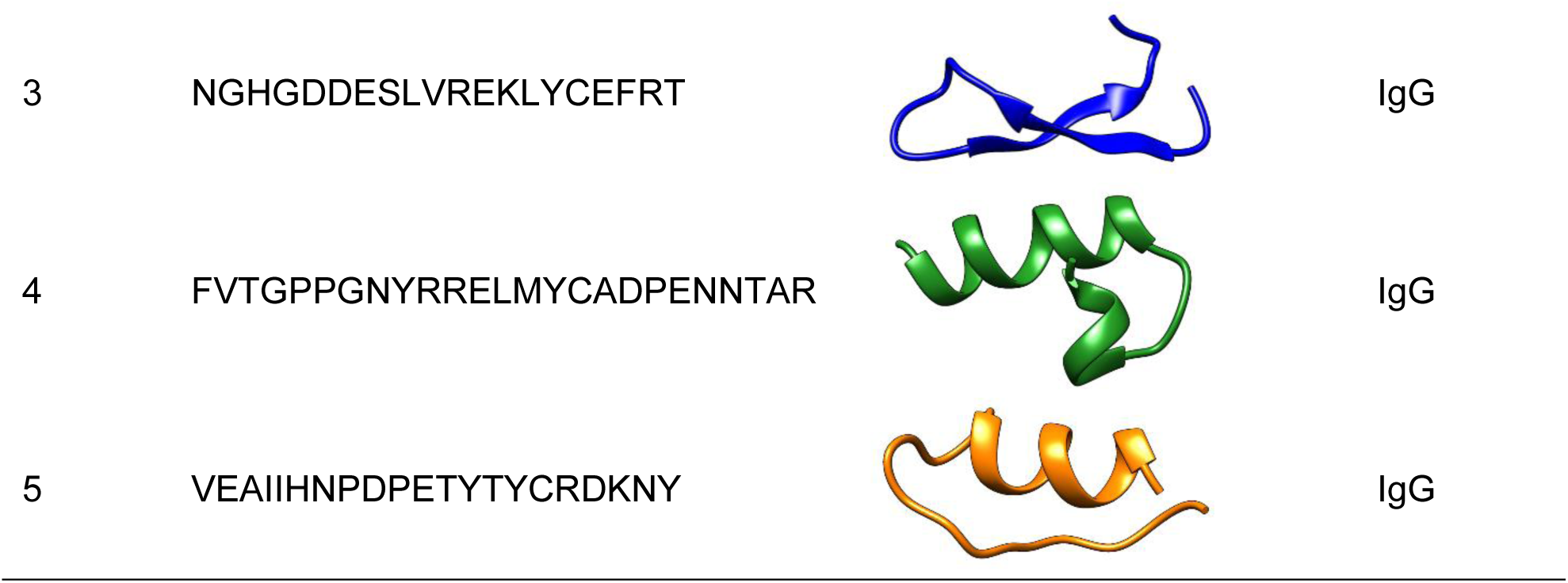
*Trypanosoma cruzi* epitope sequences with high molecular coupling.

Furthermore, *in silico* physicochemical evaluation confirmed the structural viability of these five selected sequences (ranging from 20 to 25 amino acids) as robust diagnostic targets (Table 3). All candidates were classified as stable, exhibiting instability indices well below the degradation threshold (2.19 to 32.90) and high hydrophilicity, as denoted by negative grand average of hydropathicity (GRAVY) values (-1.200 to -0.675), alongside diverse theoretical isoelectric points (3.97 to 10.26) and aliphatic indices indicating thermostability (35.20 to 69.17). Ultimately, applying the predefined statistical threshold and integrating these optimal physicochemical profiles, the five top-scoring complexes were prioritized for subsequent molecular dynamics simulations.

**Table 3.** *In silico* physicochemical properties of selected *Trypanosoma cruzi* B-cell epitopes.

| No. Epitope | No. aminoacid | Theoretical pI | Instability index | Classified protein | Aliphatic index | Grand average of hydropathicity |
| --- | --- | --- | --- | --- | --- | --- |
| 1 | 24 | 3.97 | 12.05 | stable | 36.25 | -1.200 |
| 2 | 24 | 10.26 | 27.12 | stable | 69.17 | -0.675 |
| 3 | 20 | 4.90 | 2.19 | stable | 53.50 | -1.180 |
| 4 | 25 | 6.17 | 32.90 | stable | 35.20 | -1.012 |
| 5 | 21 | 4.75 | 21.90 | stable | 55.71 | -1.124 |

The molecular dynamics simulations of the protein-peptide complexes revealed several key insights into the structural stability and interactions of the complexes over time (Fig. 5). RMSD values indicated that the complexes exhibited relatively stable conformations throughout the 500 ns simulation. Specifically, the RMSD values for the protein-peptide complexes fluctuated within a narrow range, suggesting a stable binding mode between the peptide and the protein. The RMSD values of 1.33 Å (0.3-2.4 Å), 2.44 Å (0.4-1.8 Å), 1.50 Å (0.3-3.1 Å), 1.27 Å (0.6-2.7 Å), and 1.30 Å (0.4 – 3.6 Å), corresponding to the same order of epitopes presented in Table 4, demonstrate minimal deviation from the initial conformation, reflecting the stability of the interactions throughout the simulation.

**Fig. 5.**
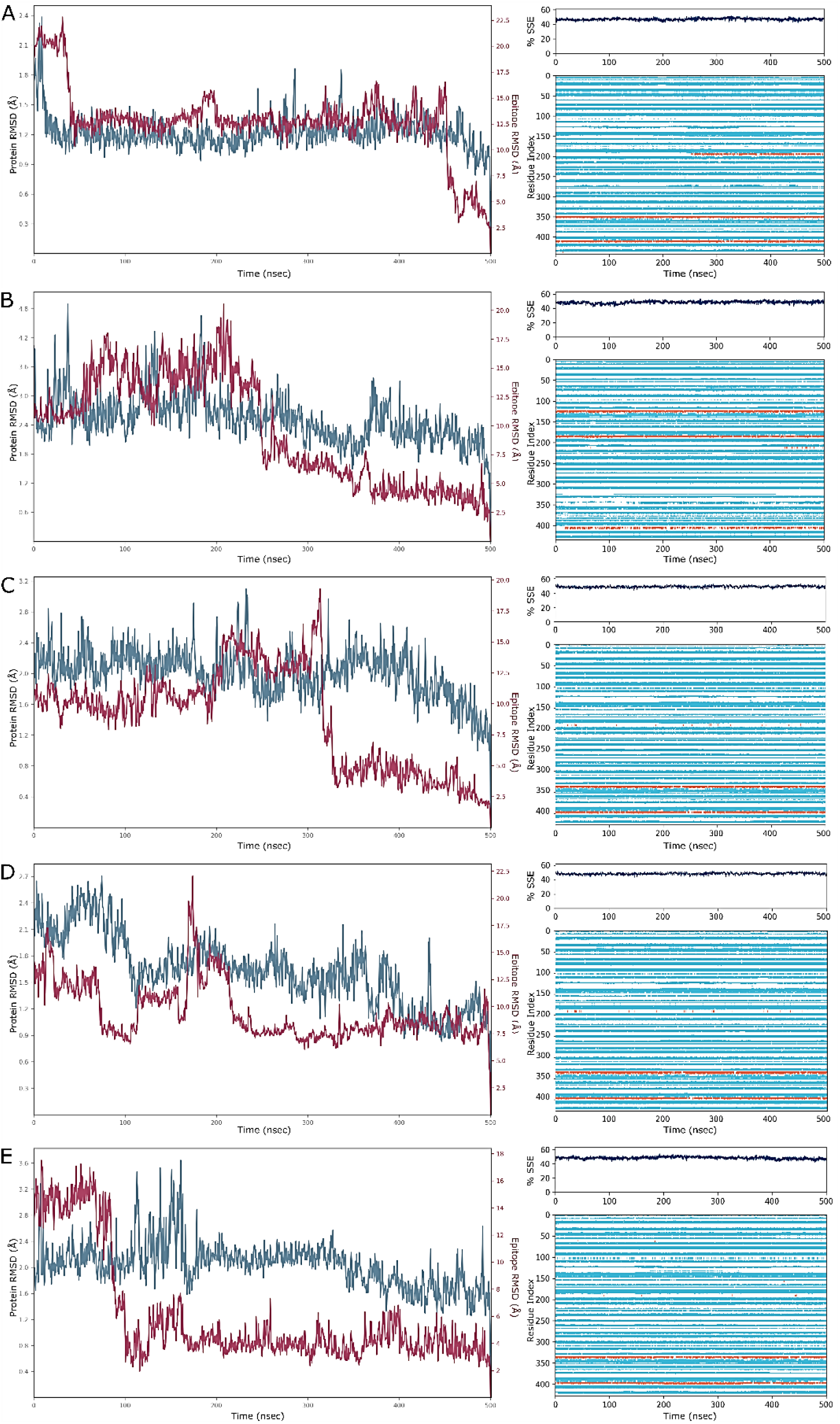
Molecular dynamics simulations of protein-peptide complexes over a 500 ns period were performed to assess their stability. The graphs (A–E, corresponding to epitopes in Table 2) display the RMSD and %SSE throughout the simulation.

**Table 4.** Diagnostic performance evaluation of epitope against patient serum samples.

| Antigens | Groups | Fisher's test | Se | 95%CI | Sp | 95%CI | Likelihood ratio |
| --- | --- | --- | --- | --- | --- | --- | --- |
| Epitope 4 | CD x HC | <0.0001 | 100 | 95.82 - 100.0 | 96.67 | 83.33 - 99.83 | 30.0 |
|  | CD x CRD | <0.0001 | 100 | 95.82 - 100.0 | 70.91 | 57.86 - 81.23 | 3.4 |
| Epitope 5 | CD x HC | <0.0001 | 100 | 95.82 - 100.0 | 96.67 | 83.33 - 99.83 | 30.0 |
|  | CD x CRD | <0.0001 | 100 | 95.82 - 100.0 | 90.91 | 80.42 - 96.05 | 11.0 |

Healthy controls (HC), cross-reactive diseases (CRD), Chagas disease (CD), Sensitivity (Se), and Specificity (Sp).

In addition, %SSE was calculated to assess the overall rigidity and stability of the complexes. %SSE values showed that the protein-peptide complexes maintained a high degree of structural integrity during the simulation, with slight fluctuations occurring only in the later stages. This indicates that the complexes remained relatively rigid, with key interactions preserved over time. Residue analysis further supported the stability findings. The analysis showed that certain residues within the protein and peptide contributed significantly to maintaining the stability of the complex, as reflected by the consistently low RMSD values and high %SSE values. These results highlight the potential of the identified peptide targets for further development as diagnostic biomarkers for Chagas disease, showing promising stability and interaction profiles in molecular dynamics simulations. The stable conformations and minimal fluctuations observed throughout the simulations suggest that the complexes may serve as reliable candidates for future diagnostic applications.

### Proof-of-concept IgG-ELISA

#### Preliminary screening and selection of candidate epitopes

To bridge the *in silico* predictions with in vitro diagnostic utility, a preliminary serological screening was conducted using a restricted panel of well-characterized, highly reactive CD-positive sera and healthy controls. Although all five synthesized peptides exhibited optimal physicochemical profiles, initial ELISA evaluations revealed that epitopes 1, 2, and 3 displayed suboptimal immunoreactivity or elevated background noise (non-specific binding) with negative sera, limiting their discriminatory capacity. This discrepancy between computational stability and *in vitro* recognition highlights the complex nature of natural antibody repertoires and peptide presentation on solid phases. Consequently, these three candidates were excluded from large-scale validation. Epitopes 4 and 5, which demonstrated robust antigenic recognition and high signal-to-noise ratios during this preliminary phase, were advanced as the definitive candidates for comprehensive serological assessment.

#### Seropositivity rates of candidate epitopes

The diagnostic potential of epitopes 4 and 5 in distinguishing Chagas disease (CD) from cross-reactive diseases (CRD) and healthy controls (HC) was evaluated using an enzyme-linked immunosorbent assay (ELISA). Sensitivity, specificity, and likelihood ratios were assessed, as summarized in Table 4 and depicted in Fig. 6. The diagnostic performance of epitope 4 showed 100% sensitivity across all tested groups (CD vs HC, CD vs CRD), with 95% confidence intervals (CI) ranging from 95.82% to 100%. Specificity for epitope 4 was notably high, with values of 96.67% for both the CD vs HC group. However, specificity for the CD vs CRD group was lower at 70.91%, with a 95% CI ranging from 57.86% to 99.83%. Fisher’s exact test demonstrated significant differences between the groups (p < 0.0001). The likelihood ratio for epitope 4 was 30.0 for the CD vs HC group, suggesting strong diagnostic utility, especially in distinguishing CD from healthy individuals. For the CD vs CRD group, the likelihood ratio was 3.4, indicating moderate utility in detecting cross-reactivity with other diseases. Similarly, epitope 5 displayed 100% sensitivity in all tested groups (CD vs HC, CD vs CRD), with 95% CI ranging from 95.82% to 100%. Specificity for epitope 5 was also 96.67% for the CD vs HC group, with a slightly higher specificity of 90.91% for the CD vs CRD group, and the 95% CI ranged from 80.42% to 96.05%. Fisher’s exact test indicated highly significant differences across all groups (p < 0.0001), reinforcing the potential of epitope 5 as a reliable diagnostic marker. The likelihood ratio for epitope 5 was 30.0 in the CD vs HC group, and 11.0 in the CD vs CRD group, further supporting its strong diagnostic value in Chagas disease detection.

**Fig. 6.**
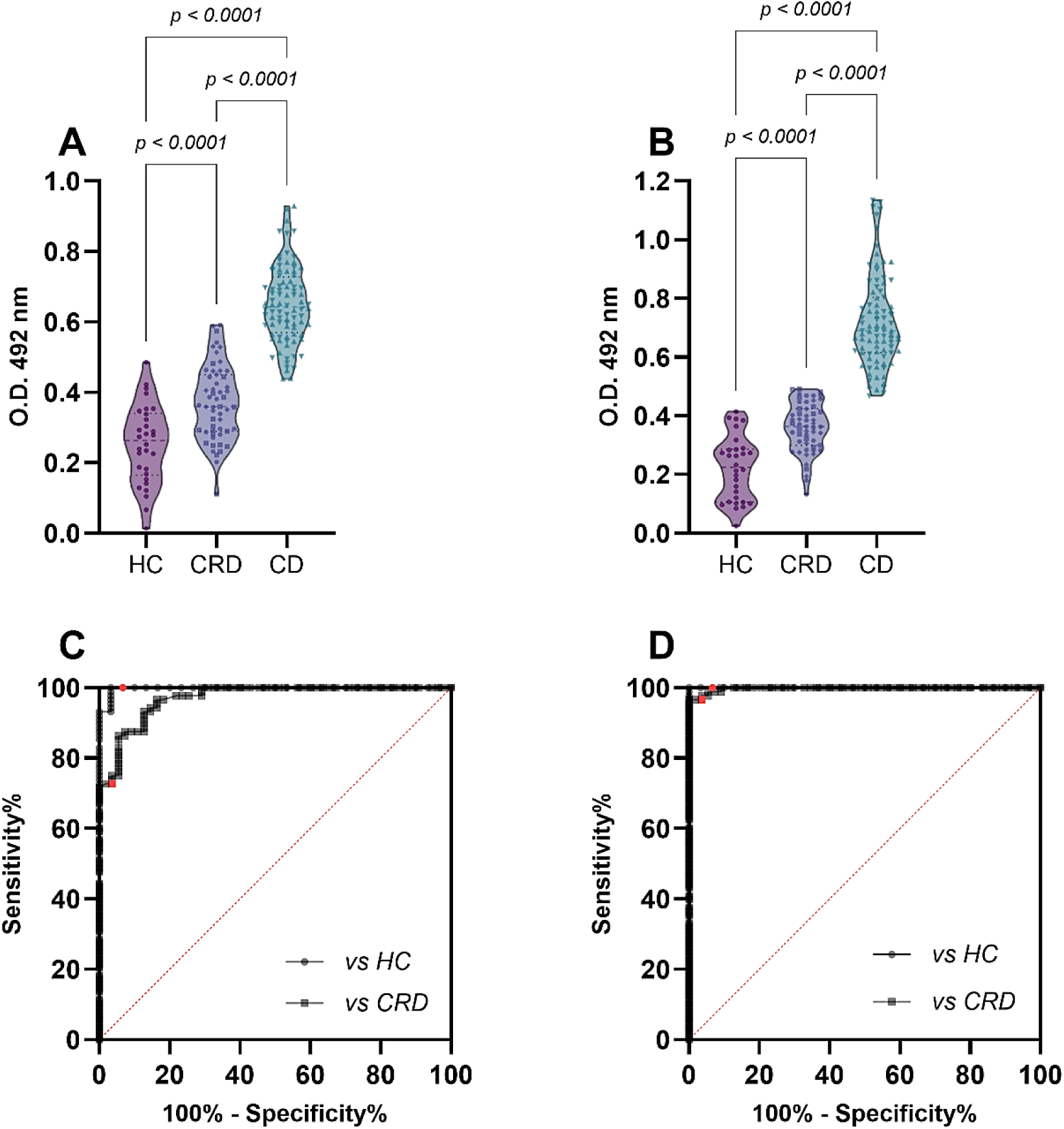
Diagnostic evaluation of epitopes 4 and 5 for Chagas disease detection. (A-B) Violin plots showing the optical density values at 492 nm obtained from ELISA for epitopes 4 and 5 across different groups: healthy controls (HC), cross-reactive diseases (CRD), and Chagas disease (CD). The significant differences between the groups are indicated by p-values < 0.0001. (C-D) Receiver operating characteristic (ROC) curves for epitopes 4 and 5, showing the sensitivity (%), specificity (%), and likelihood ratios for differentiating CD from HC and CRD.

## Discussion

This study successfully implemented an integrated computational and experimental pipeline for the rational identification and validation of *T. cruzi*-specific linear B-cell epitopes with high diagnostic potential for Chagas disease. The comprehensive approach, spanning from proteome-wide bioinformatic screening to serological confirmation, demonstrates the feasibility of this multidisciplinary strategy and reveals critical insights into the nature of *T. cruzi*-specific immunogens and their potential to address long-standing challenges in Chagas disease diagnostics.

The initial *in silico* screening phase was essential for navigating the complexity of the *T. cruzi* proteome, which is characterized by its size, redundancy, and abundance of truncated proteins. Approximately 16% of the proteome was found to consist of truncated fragments, consistent with the fragmented state of the *T. cruzi* genome assembly and highlighting the importance of quality filtering before epitope prediction (65,66). The subsequent exclusion of proteins with significant homology to human or *Leishmania* proteomes represented a critical strategic step, distinguishing this work from earlier antigen discovery efforts. This stringent filtering, which eliminated over 4,400 proteins, aimed to minimize the risk of autoimmune reactions and cross-reactivity, the principal limitations of current serological tests based on crude lysates or purified native antigens (67,68). The resulting pool of parasite-exclusive proteins provided an ideal foundation for identifying highly specific diagnostic targets.

The computational prediction of B-cell epitopes remains challenging due to the lack of a universal gold-standard algorithm. The significant variability in the number of epitopes predicted by the five different servers (APP, BepiPred 2.0, BCPred, ABCPred, and FBCPred) reflects fundamental differences in their underlying algorithms—ranging from propensity scales and machine learning to structural propensity indicators (69,70). Rather than viewing this discrepancy as a limitation, the study leveraged it to implement a consensus strategy, increasingly recognized as a best practice in computational immunology (38,71). By requiring agreement across multiple prediction tools, confidence in the candidate epitopes was significantly increased. The stratification of predictions based on the level of agreement created a valuable confidence gradient, with the 401 epitopes identified by all five tools representing the highest-confidence candidates with minimal likelihood of being false positives.

An innovative aspect of the pipeline was the incorporation of structural stability assessment early in the selection process. While many in silico studies focus solely on linear immunogenicity, it was recognized that a synthetic peptide must maintain a stable conformation in solution to function effectively in diagnostic assays (26,72). The application of the instability index filter eliminated over half of the top consensus epitopes, highlighting that immunogenic potential alone is insufficient for predicting diagnostic utility. This finding aligns with the growing recognition that intrinsically disordered peptides often fail to mimic native epitopes and perform poorly in serological assays (73). Structural modeling with PEP-FOLD 3.5 provided further confirmation that the selected epitopes possessed compact, well-folded structures compatible with antibody recognition.

The structural characterization of human Fab paratopes across different isotypes revealed important insights into potential binding interactions. The observed differences in CDR profiles between IgA, IgG, and IgM isotypes corroborate established knowledge about structural variations in antibody binding sites (74), while the predominance of heavy chain interactions aligns with the well-documented central role of heavy chain CDR3 regions in antigen recognition and specificity determination (75). These structural analyses provided a rational basis for expecting isotype-specific binding patterns in subsequent docking experiments. The molecular docking analysis revealed thermodynamically favorable binding affinities for the selected epitope-isotype pairs, yielding energy scores that significantly surpassed statistical thresholds. This differential *in silico* affinity profile—identifying distinct interactions for IgA (1), IgM (1), and predominantly IgG (3)—provides a robust structural framework for interpreting the complex humoral response observed in Chagas disease. Crucially, the translational rationale linking our multi-isotype modeling to an IgG-focused experimental validation is grounded in the clinical kinetics of Trypanosoma cruzi infection; while computational docking explored the full landscape of antibody interactions to characterize the comprehensive immunogenic potential of the epitopes, our wet-lab validation strictly prioritized IgG detection, as persistently elevated IgG titers constitute the serological gold standard for diagnosis during the chronic phase (76). Consequently, whereas the prediction of IgM and IgA interactions maps the theoretical breadth of epitope surface accessibility, the experimental corroboration of IgG confirms practical diagnostic utility in the chronic setting, a conclusion further reinforced by the visualization of dense interaction networks—stabilized by hydrogen bonds and hydrophobic contacts—within the paratope cavities (77).

The molecular dynamics simulations substantially strengthened confidence in these interactions by demonstrating their stability over biologically relevant timescales. The consistently low RMSD values (1.3-2.4 Å) throughout the 500 ns simulations indicated that the complexes maintained stable binding modes with minimal deviation from their initial conformation—a key indicator of biological relevance (78,79). The preservation of secondary structure elements (%SSE) further supported the conclusion that these epitopes form stable complexes with their cognate antibodies, meeting a critical requirement for diagnostic applications where stable antigen-antibody interactions are essential for assay reliability.

The transition from *in silico* prediction to experimental validation represented a significant challenge in computational immunology, yet the successful synthesis and purification of all five candidate peptides with high yield and purity (>95%) demonstrated the practical feasibility of the approach. The serological performance of epitopes 4 and 5 was exceptional, achieving perfect sensitivity (100%) across all patient groups—a rare achievement that surpasses most previously reported Chagas disease biomarkers (5,67). This performance is particularly notable when compared to commercial ELISA kits and chemiluminescent assays that rely on crude lysates or recombinant antigens, which often exhibit batch-to-batch variability and inconclusive results in “grey zone” patients (80). Crucially, the high specificity against both healthy controls (96.67%) and patients with cross-reactive diseases (90.91%) demonstrated specifically by epitope 5 addresses the longstanding challenge of distinguishing *T. cruzi* from Leishmania spp. in endemic regions (81,82). Unlike traditional biological antigens, the strictly defined chemical structure and superior thermal stability of our synthetic peptides position them as ideal candidates for lateral flow immunochromatographic assays, offering a robust alternative for Point-of-Care testing in rural areas where laboratory infrastructure is limited (83).

The transition from *in silico* prediction to experimental validation represented a significant challenge in computational immunology, and the successful synthesis and purification of all five candidate peptides with high yield and purity (>95%) demonstrated the practical feasibility of the approach. The serological performance of epitopes 4 and 5 was exceptional, with perfect sensitivity (100%) across all patient groups, a rare achievement that surpasses most previously reported Chagas disease biomarkers (5,67). Even more notably, the high specificity against both healthy controls (96.67%) and patients with cross-reactive diseases (90.91%) represents a potentially transformative advance. This specificity is particularly significant given the longstanding challenge of serological cross-reactivity between *T. cruzi* and other pathogens, especially *Leishmania* spp., in endemic regions (81,82).

To understand the biological basis of this exceptional immunoreactivity, we examined the parent proteins of our top-performing candidates. Epitope 4 is derived from a putative glycosyl transferase-like protein (UniProt ID: Q4D163), an enzyme class intimately associated with the metabolic processes and the synthesis of surface glycoconjugates that characterize the parasite’s interface with the host (65). Given the annotated membrane association of this protein, it is biologically plausible that this epitope represents an accessible surface motif, facilitating the robust antibody recognition observed in our study. Conversely, Epitope 5 originates from an uncharacterized protein (UniProt ID: Q4DP55) (65), a finding that underscores the strategic value of our unbiased, proteome-wide screening. By successfully mining a highly diagnostic marker from the unannotated “dark matter” of the *T. cruzi* genome, this result validates the pipeline’s capacity to identify novel immunogens that would be missed by traditional approaches focused solely on known, historically immunodominant antigens (84).

The statistical measures of diagnostic performance further underscore the potential utility of these epitopes. The positive likelihood ratios of 30.0 for both epitopes 4 and 5 against healthy controls, alongside 3.4 for epitope 4 and 11.0 for epitope 5 against cross-reactive syndrome patients, indicate that positive test results based on these epitopes would generate large and conclusive shifts in pre-test probability (85,86). These values far exceed the threshold of 10 that is typically considered indicative of a highly informative diagnostic test.

The statistical measures of diagnostic performance further underscore the potential utility of these rationally designed epitopes. The positive likelihood ratios of 30.0 for both epitopes 4 and 5 against healthy controls indicate a robust baseline specificity. Crucially, when segregating the cross-reactive cohort by specific underlying infections, the epitopes maintained high discriminatory power. Overcoming cross-reactivity with closely related trypanosomatids, particularly *Leishmania* spp., remains the primary bottleneck in Chagas disease immunodiagnostics due to extensive phylogenetic and antigenic homology (87,88). By achieving sustained positive likelihood ratios (3.4 for epitope 4 and 11.0 for epitope 5) against defined cross-reactive subgroups, our findings indicate that the *in silico* selection of these epitopes successfully evades conserved pan-trypanosomatid regions. Consequently, positive test results based on these epitopes generate large and conclusive shifts in pre-test probability (85,86), with epitope 5 exceeding the threshold of 10 typically indicative of a highly informative diagnostic test. Furthermore, the consistent recognition of these epitopes across distinct clinical forms of Chagas disease highlights their universal diagnostic coverage, regardless of the patient’s clinical staging.

When contextualized within the existing literature, our findings represent a significant paradigm shift in Chagas disease diagnostics, moving from empirical antigen selection to high-throughput rational design. Historically, diagnostic efforts have relied on a restricted set of immunodominant proteins—such as Recombinant antigen B13, Cruzain Related Antigen, or Flagellar Repetitive Antigen—which, despite their utility, frequently compromise specificity due to shared epitopes with co-endemic kinetoplastids like *Leishmania* spp. and *Trypanosoma rangeli* (16,89,90); however, by implementing a proteome-wide scanning strategy augmented by stringent homology filtering and multi-tiered computational validation, we successfully circumvented these limitations to identify novel B cell epitopes phylogenetically unique to *T. cruzi* (91,92). These candidates not only offer the requisite stability to anchor a new generation of high-precision point-of-care assays critically needed in resource-limited settings (93) but also enable the engineering of multi-epitope fusion proteins that enhance sensitivity without sacrificing established specificity (94,95).

Nevertheless, despite these promising results, this study presents limitations that define the roadmap for future validation. While the current sample size (n=178) was sufficient for proof-of-concept analysis, rigorous assessment in larger, multi-centric cohorts is necessary. Specifically, our next phase of validation involves recruiting a cohort from the southern region of Peru (e.g., Arequipa) and the Gran Chaco region. This geographic expansion is critical to evaluate the diagnostic performance of our epitopes against different *T. cruzi* Discrete Typing Units (DTUs). Since the genetic diversity of *T. cruzi* varies significantly by geography (e.g., TcI vs. TcII/TcV/TcVI), testing sera from these distinct epidemiological scenarios will ensure that the identified epitopes are universally conserved and not limited to a specific strain circulating in the initial study population (96,97). Furthermore, future studies must include sera from *T. rangeli*-infected patients to strictly rule out intrageneric cross-reactions, complementing our current Leishmaniasis and Leprosy controls, as well as evaluating these epitopes in acute-phase sera to delineate their full diagnostic utility.

## Conclusions

In conclusion, this study successfully demonstrates the power of an integrated in silico and experimental pipeline for the rational design of highly specific diagnostic antigens for Chagas disease. By implementing a stringent bioinformatic strategy that excluded proteins with homology to *Leishmania* spp. and *H. sapiens*, followed by multi-algorithm consensus epitope prediction and structural stability filtering, we identified novel *T. cruzi*-specific B-cell epitopes. The exceptional serological performance of the lead candidates, achieving 100% sensitivity and over 90% specificity, even against cross-reactive diseases, validates the central hypothesis and underscores the critical importance of reducing homology to eliminate false positives. This work not only provides a robust foundation for the development of a transformative next-generation immunoassay but also establishes a reproducible blueprint for targeted epitope discovery in complex pathogens, offering a promising path toward improved epidemiological surveillance and disease management in endemic regions.

## Institutional Review Board Statement

The study was conducted in strict accordance with the Declaration of Helsinki and approved by the Ethics Committee of the Federal University of Minas Gerais (UFMG, Belo Horizonte, Minas Gerais, Brazil). The research protocol was logged and authorized under the reference number CAAE-32343114.9.0000.5149, validating the ethical compliance of the procedures, including the collection of venous blood samples and their subsequent processing and storage at -70°C for analytical use.

## Informed Consent Statement

Informed consent was obtained from all subjects involved in the study at the time of inclusion. The cohort comprised 178 adult participants, including patients diagnosed with Chagas disease (n=88), Leishmaniasis (n=20), and Leprosy (n=20), as well as healthy controls (n=30) residing in an endemic region; all individuals provided written agreement authorizing the utilization of their biological samples and clinical data for research purposes.

## Data Availability Statement

Not applicable.

## Acknowledgments

The authors would like to thank the Coordenação de Aperfeiçoamento de Pessoal de Nível Superior (CAPES, Finance Code 001), Conselho Nacional de Desenvolvimento Científico e Tecnológico, CNPq (140491/2023-6), and Fundação de Amparo à Pesquisa do Estado de Minas Gerais, FAPEMIG (APQ-0270423, APQ-05001-22, BPD-00647 22, RED-0006723, RED-0019323). RCG, RAMA, EAFC, MSSA, and ASG would like to thank CNPq for their PQ/DT fellowship.

